# Native SAD Phasing at Room Temperature

**DOI:** 10.1101/2021.12.13.472485

**Authors:** Jack B. Greisman, Kevin M. Dalton, Candice J. Sheehan, Margaret A. Klureza, Doeke R. Hekstra

## Abstract

Single-wavelength anomalous diffraction (SAD) is a routine method for overcoming the phase problem when solving a new macromolecular structure. This technique requires the accurate measurement of intensities to sensitively determine differences across Bijvoet pairs, making it a stringent test for the reliability of a data collection method. Although SAD experiments are commonly conducted at cryogenic temperatures to mitigate the effects of radiation damage, such temperatures can alter the conformational ensemble of the protein crystal and may impede the merging of data from multiple crystals due to non-uniform freezing. Here, we propose a data collection strategy to obtain high-quality data from room temperature samples. To illustrate the strengths of this approach we use native SAD phasing at 6.5 keV to solve four structures of three model systems at 295 K. The resulting datasets allow for automatic phasing and model building, and exhibit alternate conformations that are well-supported by the electron density. The high-redundancy data collection method demonstrated here enables the routine collection of high-quality, room-temperature diffraction to improve the study of protein conformational ensembles.

## 1 Introduction

X-ray crystallography is the predominant experimental technique for determining the structures of macromolecules at atomic resolution. This method reconstructs the electron density of a crystal from measurements of the relative intensities of scattered X-rays, enabling researchers to build a model with atomic detail. Although the intensities of the scattered X-rays determine the amplitude of the diffracted waves, the phases of those waves are needed to reconstruct the electron density [1]. For the structures of novel macromolecules, this problem is often overcome using the resonance phenomenon of anomalous scattering, with single-wavelength anomalous diffraction (SAD) being the method of choice for the *de novo* phasing of macromolecular structures [2, 3].

SAD phasing relies on the accurate measurements of the differences between Bijvoet pairs of reflections. An especially challenging case is native SAD phasing, which relies on the weak anomalous scattering from low-Z atoms of the macromolecule or the crystallization conditions. Although the first *de novo* SAD structure used the anomalous scattering from native sulfur atoms [4], structures solved by native SAD phasing still constitute a small fraction of those deposited in the Protein Data Bank [5]. Because of its challenge, native SAD phasing is commonly used as a proving ground for new technologies and data collection methods [6, 7, 8, 9].

High-redundancy data collection strategies improve the accurate determination of Bijvoet differences and therefore increase the chances of successful phasing by the SAD method [10, 6, 8]. However, cryogenic temperatures are typically used to mitigate the resulting risk of radiation damage [11, 12], and the cryocooling of crystals can alter the ensemble of conformations [13] and can limit the merging of multiple crystals due to imperfect isomorphism [14]. Since the first successful S-SAD experiment [4], native SAD experiments have been reported nearly exclusively under cryogenic conditions (with the exception of serial crystallography applications [7]).

Room-temperature X-ray crystallography is seeing a resurgence to better probe the dynamics of protein structures at physiologically relevant temperatures [15, 16, 17]. However, a common concern with such experiments is the possibility of radiation damage impacting the results. Here, we demonstrate native SAD phasing from single crystals at room temperature. We present four structures of three different model enzymes, which we solved by automatic SAD phasing and model building, and exhibit alternate conformations that reflect the conformational heterogeneity of the crystals. Our high-redundancy data collection strategy demonstrates the accuracy of room-temperature diffraction experiments and can be implemented at many macromolecular beamlines.

## 2 Materials and methods

### 2.1 Protein Purification and Crystallization

#### 2.1.1 Hen Egg-White Lysozyme

We dissolved lyophilized hen egg-white lysozyme (Sigma-Aldrich), hereafter HEWL, in deionized water to concentrations of 15-30 mg/mL. Using the hanging drop vapor diffusion method, we grew tetragonal lysozyme crystals with well solution composed of 50 mM sodium citrate buffer (pH 5.2-5.8), 20-25% glycerol, and 20-25% PEG4000. We set drops with 2 µL protein solution and 2 µL well solution over 0.5 mL wells and stored the crystallization plates at ambient temperature (approximately 22°C). Tetragonal lysozyme crystals (0.2-0.5 mm) grew after 3-7 days.

#### 2.1.2 *E. coli* Dihydrofolate Reductase

We expressed wildtype *E. coli* dihydrofolate reductase, hereafter DHFR, from a pET22b vector generously provided by the lab of James Fraser (UCSF) [13, 18]. We transformed BL21(DE3) *E. coli* cells (Novagen) with the plasmid and grew overnight cultures in lysogeny broth with ampicillin. We diluted the overnight cultures 1:100 in 2 L of terrific broth with ampicillin, which we grew to an optical density of 0.6-0.8 at 600 nm. Once the cultures reached the mid-log phase, we added isopropyl *β*-D-1-thiogalactopyranoside (IPTG, Invitrogen) to a final concentration of 1 mM, inducing DHFR expression. After 4-8 hours of expression, we harvested the cells by centrifugation, flash froze the resulting cell pellets in liquid nitrogen, and stored them at -80°C until purification.

We based our DHFR purification protocol on the thesis of Michael Sawaya [19]. Briefly, we thawed, resuspended, and lysed the cell pellets by sonication. We removed the cell debris by centrifugation, then used streptomycin precipitation to remove nucleotides from the supernatant. We further purified the protein fraction by ammonium sulfate precipitation at 40% saturation. The supernatant was retained and subjected to an additional ammonium sulfate precipitation at 90% saturation. The pellet was resuspended and we isolated DHFR by methotrexate-affinity chromatography using methotrexate-agarose (Sigma-Aldrich) followed by anion-exchange chromatography using a HiPrep Q FF column (Cytiva). We then pooled and concentrated the eluted protein. To minimize nucleotide contamination, we only pooled fractions that had an *A*_280_*/A*_260_ > 1.7 based on an absorbance reading with a NanoDrop spectrophotometer (Thermo Fisher Scientific). Finally, we exchanged the purified protein into 10 mM HEPES buffer (pH 7.0) containing 1 mM dithiothreitol (DTT) using an Amicon centrifugal concentrator (MilliporeSigma), flash froze aliquots in liquid nitrogen, and stored them at -80°C.

DHFR complexed with NADP^+^ and folate is commonly used as a model of the Michaelis complex of the enzyme [20]. We mixed 20 mg/mL DHFR with three-fold molar excess of NADP^+^ and folate in 20 mM imidazole (pH 7.0) with 2 mM DTT, then incubated the mixture on ice for 30 minutes. Based on the conditions of Keedy *et al* [13], we crystallized the complex using the sitting drop vapor diffusion method with a well solution composed of 20 mM imidazole (pH 5.4 - 5.8), 125 mM manganese(II) chloride, and 16-21% PEG400 (v/v). The drops contained 0.2 µL of 7-14 mg/mL protein and 0.2 µL of well solution, producing rod-shaped crystals (approximately 0.1 mm x 0.3-1 mm) after 2-4 weeks of incubation at 4°C.

#### 2.1.3 Human Protein Tyrosine Phosphatase 1B

The human protein tyrosine phosphatase 1B, hereafter PTP1B, experiments used a construct containing residues 1–321 with a C32S/C92V double mutation [21]. We expressed PTP1B from a pET24b vector with a kanamycin resistance gene, which was generously provided by the lab of James Fraser (UCSF). We expressed and purified PTP1B in accordance with published methods [22] with minor modifications. Briefly, we transformed Rosetta(DE3) *E. coli* cells (Novagen) with the PTP1B plasmid and grew overnight cultures in lysogeny broth with kanamycin and chloramphenicol. We diluted the overnight cultures 1:1000 in 2 L of terrific broth with kanamycin and chloramphenicol. We incubated the final cultures at 37°C and grew them to an optical density of 0.6 −0.8 at 600 nm. Once the cultures reached the mid-log phase, we added IPTG (Invitrogen) to a final concentration of 1 mM, inducing PTP1B expression, and incubated them at 20°C for 12-18 hours. We harvested the cells by centrifugation, flash froze the resulting cell pellets in liquid nitrogen, and stored them at -80°C until purification.

We thawed the cell pellets, resuspended them in lysis buffer, and lysed the cells by sonication. Cell debris was cleared by centrifugation and syringe filtration through a 0.45 µm filter. We purified PTP1B by cation-exchange chromatography using a HiPrep SP FF column (Cytiva). We pooled the PTP1B fractions from the elution, concentrated them using an Amicon centrifugal concentrator (MilliporeSigma), and further purified PTP1B by size-exclusion chromatography with a HiLoad Superdex 75 column (Cytiva). After this column, the purified protein was in 10 mM Tris, pH 7.5, with 25 mM NaCl and 0.2 mM EDTA. It was concentrated using an Amicon centrifugal concentrator (MilliporeSigma) and stored at -80°C until crystallization. To avoid oxidation of the catalytic Cys215 [23], we added fresh DTT (1-3 mM) to each buffer used during purification.

Prior to crystallization, we diluted PTP1B to 10-15 mg/mL using protein storage buffer. For the TCS401-bound complex, we added a 25 mM stock of TCS401 (Tocris Bioscience) dissolved in DMSO to the protein solution to a five-fold molar excess, and incubated it on ice for at least 30 minutes. We crystallized both apo and TCS401-bound PTP1B using the hanging drop vapor diffusion method with a well solution composed of 100 mM HEPES (pH 6.8-7.6) with 250 mM magnesium acetate, 11-15% PEG8000 (w/v), 10% glycerol (v/v), 6% ethanol (v/v), and 0.1% BME (v/v). Drops were set by mixing 1 µL of protein and 1 µL of well solution over 0.5 mL wells. Crystallization plates were incubated at 4°C, and rod-shaped crystals with hexagonal faces grew to 0.2-1 mm after 1-2 weeks.

### 2.2 Data Collection

We collected the datasets presented in this work at the Northeastern Collaborative Access Team (NE-CAT) beamline 24-ID-C of the Advanced Photon Source (Argonne National Laboratory) on July 17, 2019. We looped all crystals on-site using the MicroRT system (MiTeGen) for room-temperature data collection, and conducted the diffraction experiments at ambient temperature, which was measured to be approximately 295 K. We used a beam energy of 6.5 keV to maximize the native anomalous scattering contribution within the energy range of the beamline.

To obtain high-redundancy datasets while minimizing radiation damage, we collected 1440 image passes on each crystal with an exposure time of 0.1 s and an oscillation angle of 0.5 degrees. The incident X-ray intensity was attenuated to 0.5% transmission, and we collected the datasets while continually translating along the crystals to distribute the dosage across the crystal volume. The Pilatus 6M-F detector (Dectris) has 2463 × 2527 pixels, and was positioned at the minimal distance of 150 mm. Due to the long-wavelength used, the resolution was limited by the experimental geometry rather than the crystal quality for each sample. To obtain the highest resolution data accessible, we raised the detector by two panels during collection. We used this experimental geometry for both passes on HEWL, both passes on PTP1B in complex with TCS401, and both crystals of apo PTP1B. For DHFR, we collected the first pass with a centered detector and the following two passes with a raised detector to record higher resolution reflections.

### 2.3 Data Reduction, SAD Phasing, and Structure Refinement

We processed each pass on each crystal using *DIALS* [24]. Due to the raised detector geometry, we set the beam center to (290.5, 225.2) and used a pixel mask to remove a shadow cast by the beamstop support. Our images contain leakage from a higher-energy undulator harmonic. Therefore, we excluded strong reflections lower than 6 Å in resolution during indexing due to a secondary diffraction pattern originating from the harmonic. To index each pass on the PTP1B crystals, we used local index assignment (`index.assignment.method=local`) which significantly improved the percentage of indexed strong reflections. After integration, we scaled and merged the multiple passes for each crystal using *AIMLESS* [25]. For apo PTP1B, we scaled and merged one pass each from two different crystals, which we found to be necessary to improve the anomalous signal for SAD phasing. For the PTP1B:TCS401 complex, we excluded the last 360 frames from the second pass due to weaker diffraction.

We solved each structure by SAD phasing using default settings in *AutoSol* [26] *in Phenix* [27, 28]. *We used sulfur* as the anomalous scattering element for HEWL and PTP1B, and we specified manganese, phosphorous, and sulfur as the anomalous scattering elements for DHFR. We successfully phased all datasets by this approach without additional intervention, and constructed initial models for HEWL, DHFR, and TCS401-bound PTP1B using *AutoBuild* [29]. We further refined these models by cycles of automatic refinement with *phenix.refine* and manual model building in *Coot* [30]. Due to the extensive alternate conformations in the apo PTP1B electron density, we used the TCS401-bound model to initialize the apo model. We manually built the open state of the WPD loop (residues 176-193) in *Coot*, and used group occupancies in *phenix.refine* to constrain mutually-exclusive conformations to sum to full occupancy.

### 2.4 Assessing the impact of redundancy on anomalous signal

To characterize the effect of high-redundancy data collection, we scaled and merged each of the four datasets with increasing numbers of consecutive frames in *AIMLESS* [25]. To understand the impact on the precision of anomalous differences, we computed half-dataset correlation coefficients of the anomalous differences (*CC*_*anom*_) for different numbers of frames. We used *reciprocalspaceship* [31] to implement repeated two-fold cross-validation to determine the mean and standard deviations of the *CC*_*anom*_ by resolution bin for the HEWL sulfur SAD dataset.

To assess the impact on real-space anomalous signal, we computed anomalous difference maps for the datasets with different numbers of frames using the phases from the refined models. We determined anomalous peak heights using *phenix.find_peaks_and_holes* [27, 28]. For each of the datasets, we fit the anomalous peak height of each site, *s*, as a function of number of frames, *N*_*n*_, by minimizing the following least-squares objective function with the Levenberg-Marquadt algorithm implemented in *scipy* [32]:

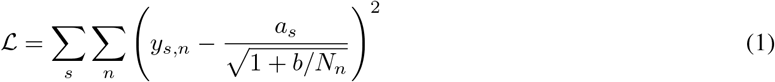

where *y*_*s,n*_ is the anomalous peak height of site *s* with *N*_*n*_ frames, *a*_*s*_ is the maximal peak height for site *s*, and *b* is a global parameter for the dataset. This functional form is derived in Appendix 8.1. Using, *a*_*s*_, we define rescaled observations as *ŷ*_*s,n*_ = 100 *y*_*s,n*_*/a*_*s*_ corresponding to the percent of the maximum attainable anomalous peak height. The rescaled data were plotted with the normalized model given by 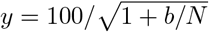, as well as a function of the form 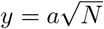 that was fit with respect to the rescaled data.

## 3 Results

### 3.1 Native SAD phasing of model systems at 295 K

Native SAD phasing is commonly used as a benchmark for the accuracy of new data collection strategies and technologies [6, 7, 8, 9]. Here, we use native SAD to solve four structures of three model systems at room temperature (295 K). Previous work by Weinert *et al* demonstrated high-redundancy data collection from single-crystal diffraction at 6 keV and at cryogenic temperatures [8, 33]. The data collection strategy presented here for native SAD phasing at room temperature differs from their approach in several ways. They used a multi-axis goniometer to collect at multiple sample orientations, while we instead continually translated along the crystals to distribute the X-ray dose across the crystal volume. In addition, Weinert *et al* used fine *ϕ*-slicing to improve their data quality. Fine *ϕ*-slicing is most effective when the rotation angle is comparable to half the crystal mosaicity [34], which, in our experience, is often not possible with room-temperature crystals due to their low crystal mosaicity (0.01-0.04°, Table 1).

**Table 1:**
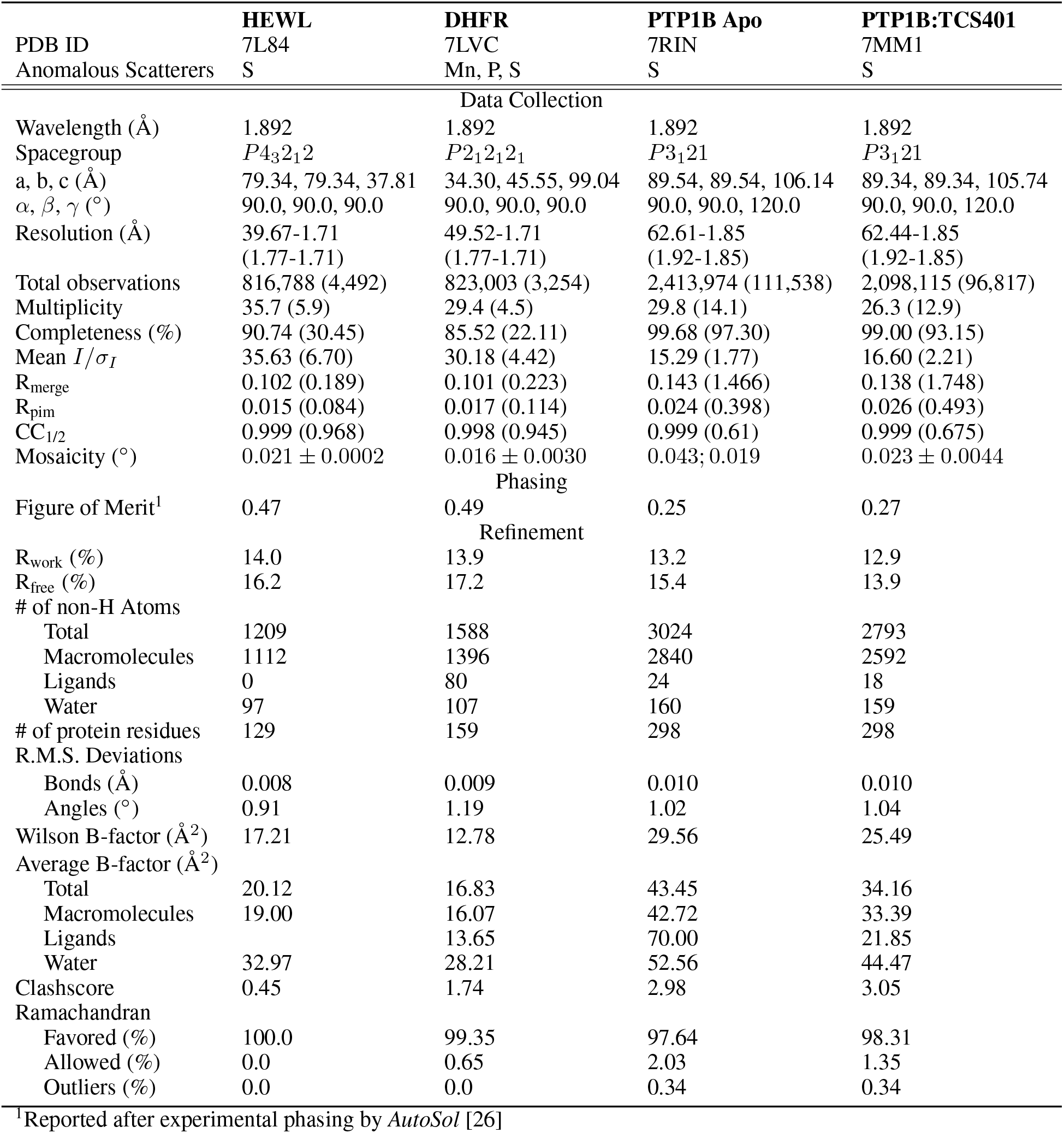
Data Collection and Refinement Statistics

Data were collected at 6.5 keV to maximize the anomalous scattering signal within the accessible spectrum of the APS 24-ID-C beamline. Each of the following structures were solved from multiple 1440-image passes on a single crystal, except for apo PTP1B which was solved from passes collected on two different crystals. The data collection statistics for these four datasets are presented in Table 1. These datasets have very high redundancy (26-35x overall), and the resolution was limited by the experimental geometry at the long wavelength leading to high mean *I/σ*_*I*_ and CC_1/2_ values and low completeness in the high resolution bins. Although the absorbed X-ray dose is higher at lower beam energies [35], the relative image B-factors fit by *AIMLESS* suggest that radiation damage did not impact data reduction [25]. Although the final passes for both DHFR and the PTP1B:TCS401 complex reach B-factors of less than -10 (Fig. S1), removing the passes did not improve SAD phasing.

The structure of HEWL was solved using the anomalous scattering from the ten sulfur atoms in the protein, and the anomalous difference map shows clear density for the four disulfide bonds and two methionine residues in the model (Fig. 1a). The Michaelis complex of DHFR was solved using the anomalous scattering from the manganese ions in the crystallization solution, the phosphorous atoms in the NADP^+^ cofactor, and the sulfur atoms in the protein. The refined model includes five ordered Mn^2+^ ions, which were supported by the anomalous difference map (Fig. 1b). Significant anomalous density (> 5*σ*) is also seen for the three phosphorous atoms in NADP^+^ and six of the seven sulfur-containing side chains. PTP1B was solved by sulfur SAD in an apo state and in complex with the active site inhibitor TCS401 [36]. Thirteen of fifteen sulfur sites were resolved in the anomalous difference map of apo PTP1B at 5*σ* (Fig. 1c), and fifteen of sixteen sulfur sites can be seen in the corresponding map of the PTP1B:TCS401 complex (Fig. 1d). In each of these structures, the missing sulfur-containing sidechains belong to methionine residues with weak density at or near the N-terminus. The refinement statistics for the model systems are summarized in Table 1.

**Figure 1:**
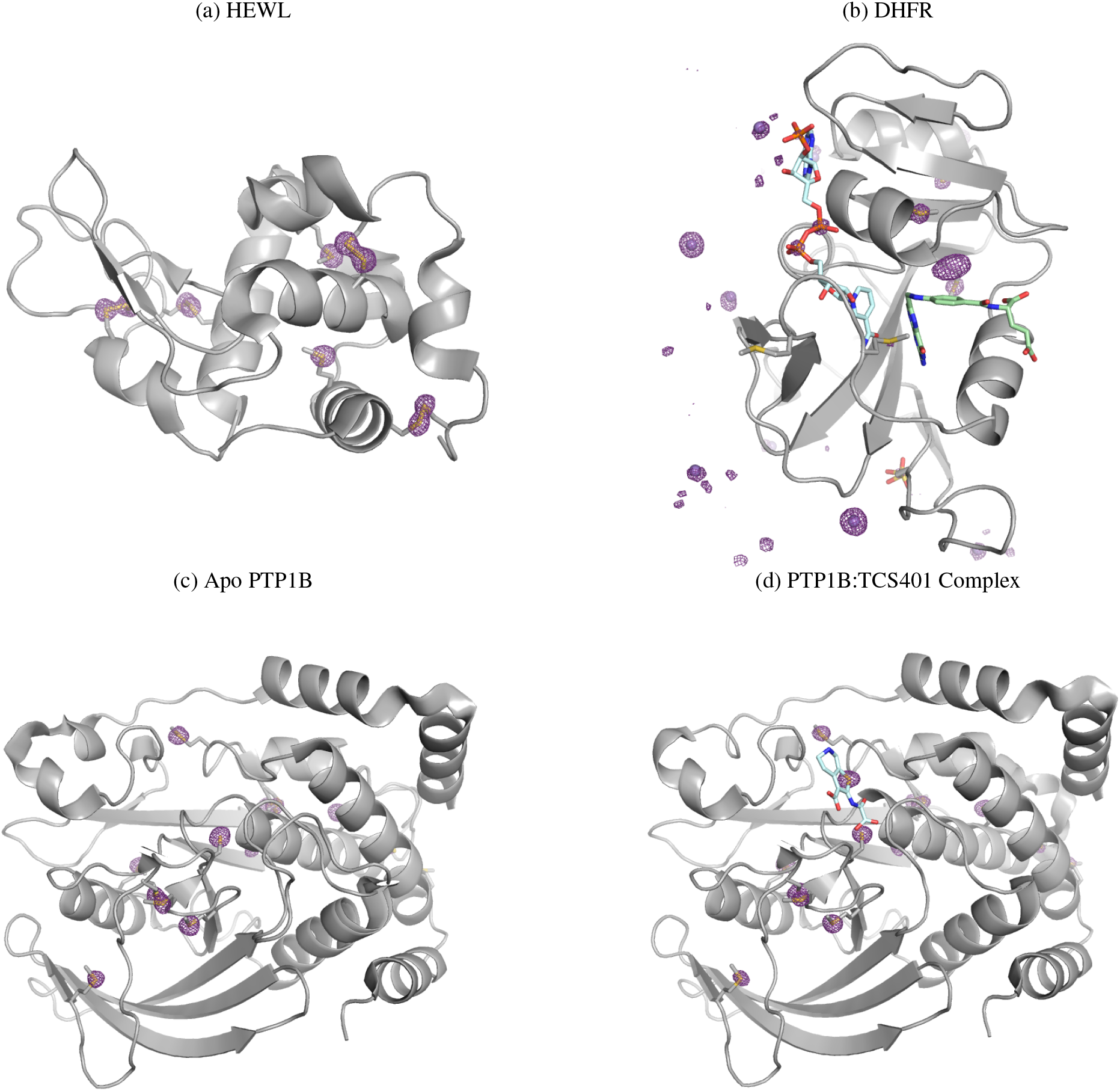
High-redundancy data collection enables room-temperature, native SAD phasing of model systems. SAD solution structures for (a) Hen egg-white lysozyme, (b) Model of the *E. coli* dihydrofolate reductase Michaelis complex, (c) Apo human protein tyrosine phosphatase 1B, and (d) TCS401-bound human protein tyrosine phosphatase 1B are shown with their respective anomalous difference maps (purple mesh). The maps in (a), (c), and (d) are contoured at 5*σ* and shown within 2 Å of modeled atoms. The DHFR anomalous difference map in (b) is contoured at 6*σ* and shown within 8 Å of modeled atoms due to the additional anomalous signal from partial-occupancy manganese sites at crystal contacts. Images were rendered using PyMOL [37].

### 3.2 DHFR anomalous differences highlight multiple anomalous scattering elements

Although the DHFR dataset was collected with the goal of SAD phasing based on the manganese content from the crystallization conditions, we observe additional anomalous difference peaks corresponding to the native sulfur atoms in the protein and the phosphorous atoms of the NADP^+^ cofactor. The active site of the refined model is shown in Figure 2a, which highlights sulfur and phosphorous anomalous peaks. In addition to the well-ordered NADP^+^ cofactor, significant anomalous differences are seen for the catalytically important Met20 and Met42 sidechains. Mutations to Met42 significantly impact hydride transfer in the enzyme [38], while Met20 shows multiple rotamers in high-resolution structures of DHFR and plays a role in both hydride transfer and regulating solvent accessibility to promote protonation of the substrate [20, 39].

**Figure 2:**
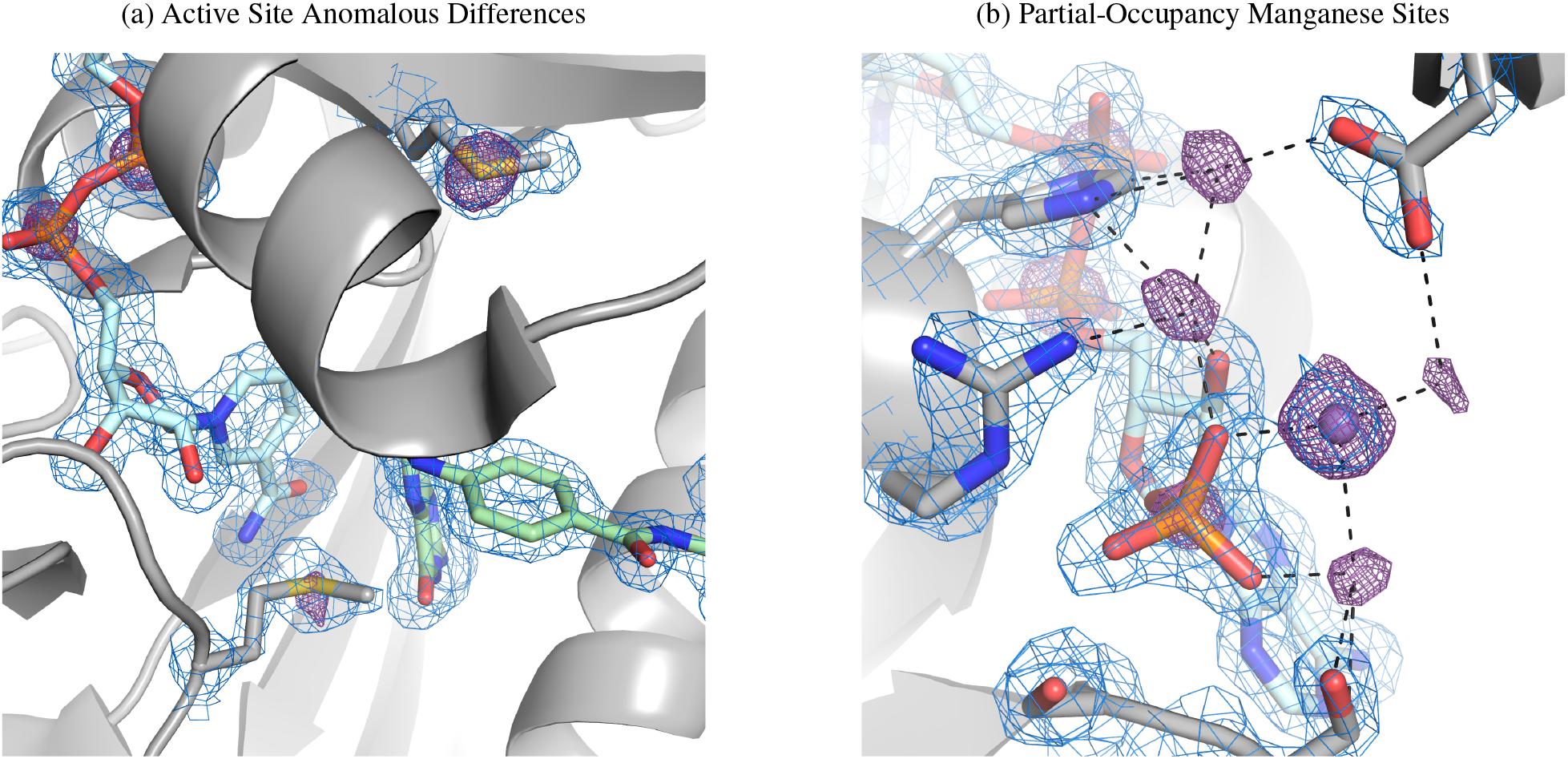
Anomalous differences in a model of the DHFR Michaelis complex highlight active-site residues and partial-occupancy ions. (a) The active site of DHFR is shown with NADP^+^ and folate, a substrate mimetic, in light-blue and green, respectively, with nearby methionine residues shown in gray. The 2*F*_*o*_ − *F*_*c*_ map is displayed in blue at 1.2*σ* within 1.5 Å of shown atoms. Density for the sulfur atoms of methionine residues and the phosphorous atoms of NADP^+^ can be seen in the anomalous difference map (purple mesh; 5*σ*). (b) Multiple partial-occupancy manganese sites are supported by the anomalous difference map (purple mesh; 5*σ*) around a phosphate group of NADP^+^. Only one of these sites was modeled as a manganese based on density in the 2*F*_*o*_ − *F*_*c*_ map (blue mesh; 1.5*σ*). The 2*F*_*o*_ − *F*_*c*_ map is displayed within 2 Å of the shown atoms, and the anomalous difference map is shown within 8 Å of the phosphorous atom. Contact distances less than 3.0 Å to manganese sites are shown as black dashed lines. Images were rendered using PyMOL [37].

The anomalous difference map also contains significant density (>5*σ*) in regions that are not supported by strong electron density in the 2*F*_*o*_ − *F*_*c*_ map (Fig. 2b). Although one manganese site is strongly supported in the 2*F*_*o*_ − *F*_*c*_ in the vicinity of the phosphate group of NADP^+^, the additional anomalous density can be best explained by partial-occupancy manganese sites around the phosphate moiety. Each of these sites involves close contacts (<3.0 Å) to protein sidechains or the phosphate group to support their coordination geometries; however, the very close distances between manganese sites (2-3 Å) suggest that occupancy at many of these sites is mutually exclusive. This underscores that the anomalous difference density represents an ensemble- and time-averaged view of the region. The close vicinity of Asp11 from a neighboring copy of DHFR also suggest that these sites are stabilized by crystal packing interactions. Observation of these partial-occupancy binding sites highlights the accuracy and sensitivity of these anomalous difference maps.

### 3.3 Room-temperature structures feature conformational heterogeneity

An advantage of room-temperature data collection is that the conformational ensemble of the macromolecule will be better represented than at cryogenic temperatures [17, 16, 13]. This is most evident in our native SAD structure of apo PTP1B, which shows a superposition of the open and closed states of the catalytically important WPD loop (Fig. 3a). Our structure refined to 61% open and 39% closed, which is consistent with the multi-temperature refinement by Keedy *et al* who obtained 65% open and 35% closed at 277 K [22]. The small difference in refined occupancy may be attributable to different refinement protocols, in particular with regard to the partial-occupancy waters in the vicinity of the WPD loop. In contrast to the apo enzyme, in the presence of TCS401, an orthosteric inhibitor, the WPD loop fully occupies the closed state (Fig. 3b). This is consistent with existing structures of PTP1B in complex with the inhibitor [36, 40].

**Figure 3:**
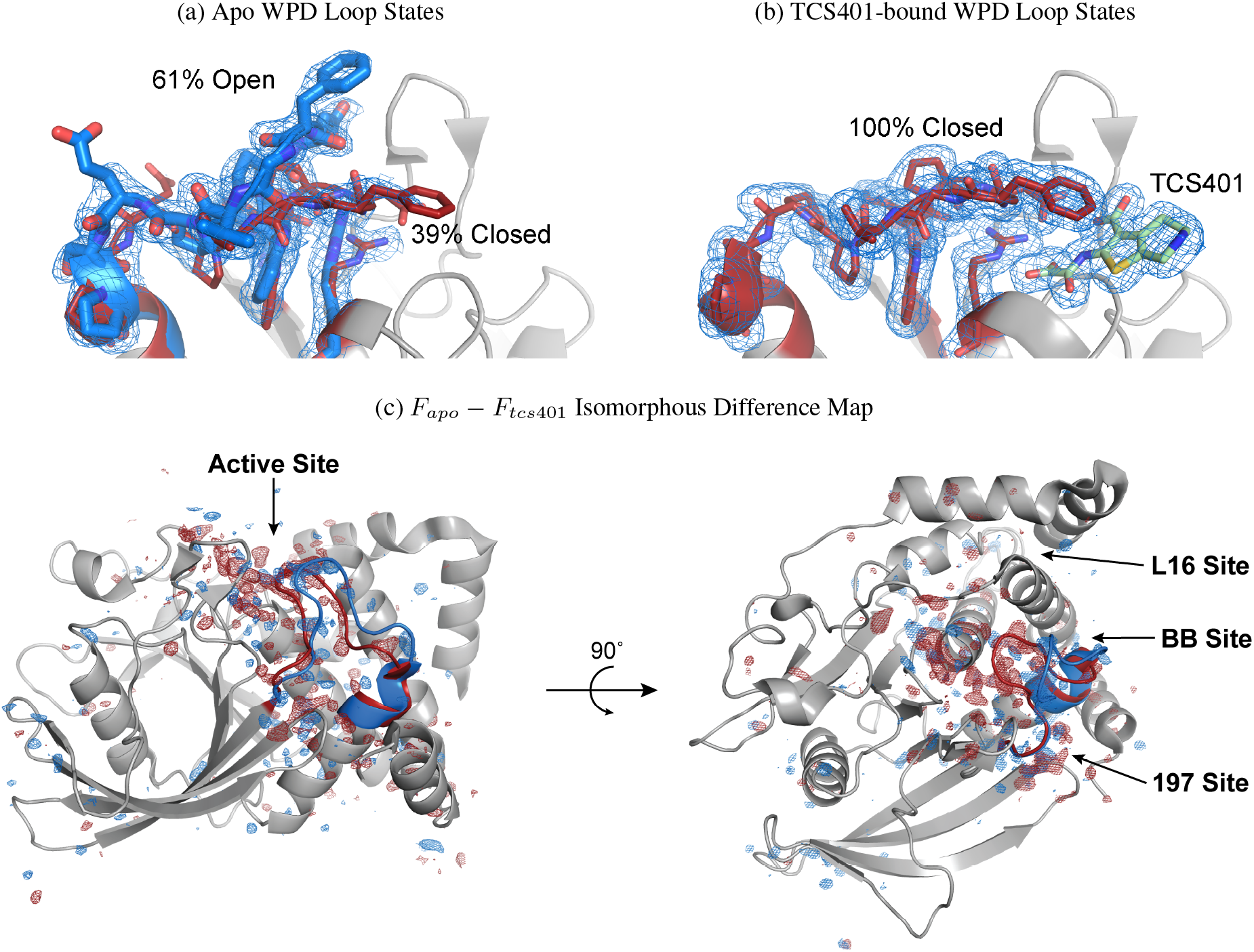
Room-temperature structures of PTP1B exhibit extensive conformational heterogeneity. (a) At room temperature, apo PTP1B adopts a superposition of the WPD open and WPD closed states which results in different positioning of important catalytic residues. (b) With a bound orthosteric inhibitor, TCS401 (green), the WPD loop is only observed in the WPD closed state. 2*F*_*o*_ − *F*_*c*_ maps are shown as blue mesh (0.5*σ*) within 1.5 Å of the displayed residues. (c) The *F*_*apo*_ − *F*_*tcs*401_ isomorphous difference map, phased using the apo model, illustrates the sites of the protein that differ on TCS401 binding. The difference map is contoured at +3.5*σ* (blue) and − 3.5*σ* (red). The WPD loop is depicted in blue (open state) and red (closed state), and known allosteric sites are labeled. Most of the structural changes that occur between WPD loop states are localized to one side of the central beta sheet of PTP1B. Images were rendered using PyMOL [37].

Because the apo structure exhibits a superposition of WPD loop states and the TCS401-bound complex fully populates the WPD closed state, we can visualize the regions that differ between the two WPD loop states using an isomorphous difference map, |*F*_*apo*_| − |*F*_*tcs*401_|. We used SCALEIT [41, 42] to place the TCS401-bound dataset on the same scale as the apo dataset, and we then computed an isomorphous difference map using the phases from the apo model. The isomorphous difference map is shown in Figure 3c overlaid on the apo structure. The positive density (blue) highlights regions that are associated with the WPD open state (stronger in the apo dataset), and the negative density (red) highlights those associated with the WPD closed state (stronger in the TCS401-bound dataset). The isomorphous difference map illustrates that the primary sites that undergo structural changes between the two WPD loop states are localized to one side of the central *β*-sheet of PTP1B. Furthermore, strong difference density is observed in the vicinity of previously identified allosteric sites such as the benzbromarone binding site (BB site) [43], L16 site [22, 44], and the 197 site [22, 44, 40]. The observation of these widespread structural differences within and between datasets illustrates a benefit of working near physiological temperatures [13], and suggests that such experiments can be used to probe allosteric mechanisms in protein structures.

### 3.4 High-redundancy data collection improves anomalous signal

To assess the impact of redundancy on anomalous signal, we reprocessed the HEWL S-SAD dataset with increasing numbers of diffraction images. To our initial surprise, anomalous peak height at sulfur atoms did not increase proportional to the square root of the number of observations, as might be expected based on a naive interpretation of models of anomalous signal [10, 2]. To account for this, we note that Terwilliger *et al*’s analysis [45] predicts a square-root dependence on the number of unique reflections, not the total number of observations. We extended their analysis to explicitly account for the redundancy of observations (Appendix 8.1). Based on our model, we expect the average anomalous peak height to be proportional to 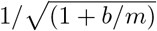, where *m* is the average multiplicity of the observed reflections and *b* is a fitting parameter which reflects data quality and the presence of background anomalous scattering by lighter elements. Because the average multiplicity of observed reflections is directly proportional to the number of frames, the average multiplicity can be equivalently expressed in terms of the number of frames, *N*.

The HEWL S-SAD dataset shows an improvement in the correlation coefficient of anomalous differences across half-datasets with increasing numbers of frames (Fig. 4a). However, the mean anomalous peak heights of the sulfur atoms approaches an asymptote with increasing numbers of frames. Our model provides an excellent fit to the anomalous peak heights, while a square-root dependence does not capture the trend seen in Fig. 4b. The fit to the anomalous peak heights yields *b* ≈750. For a dataset composed of 2878 frames, this corresponds to achieving 89% of the maximal anomalous peak height at this completeness and resolution. This model of the relationship between redundancy of observations and anomalous signal is also consistent with the three other model systems used in this work (Fig. S2).

**Figure 4:**
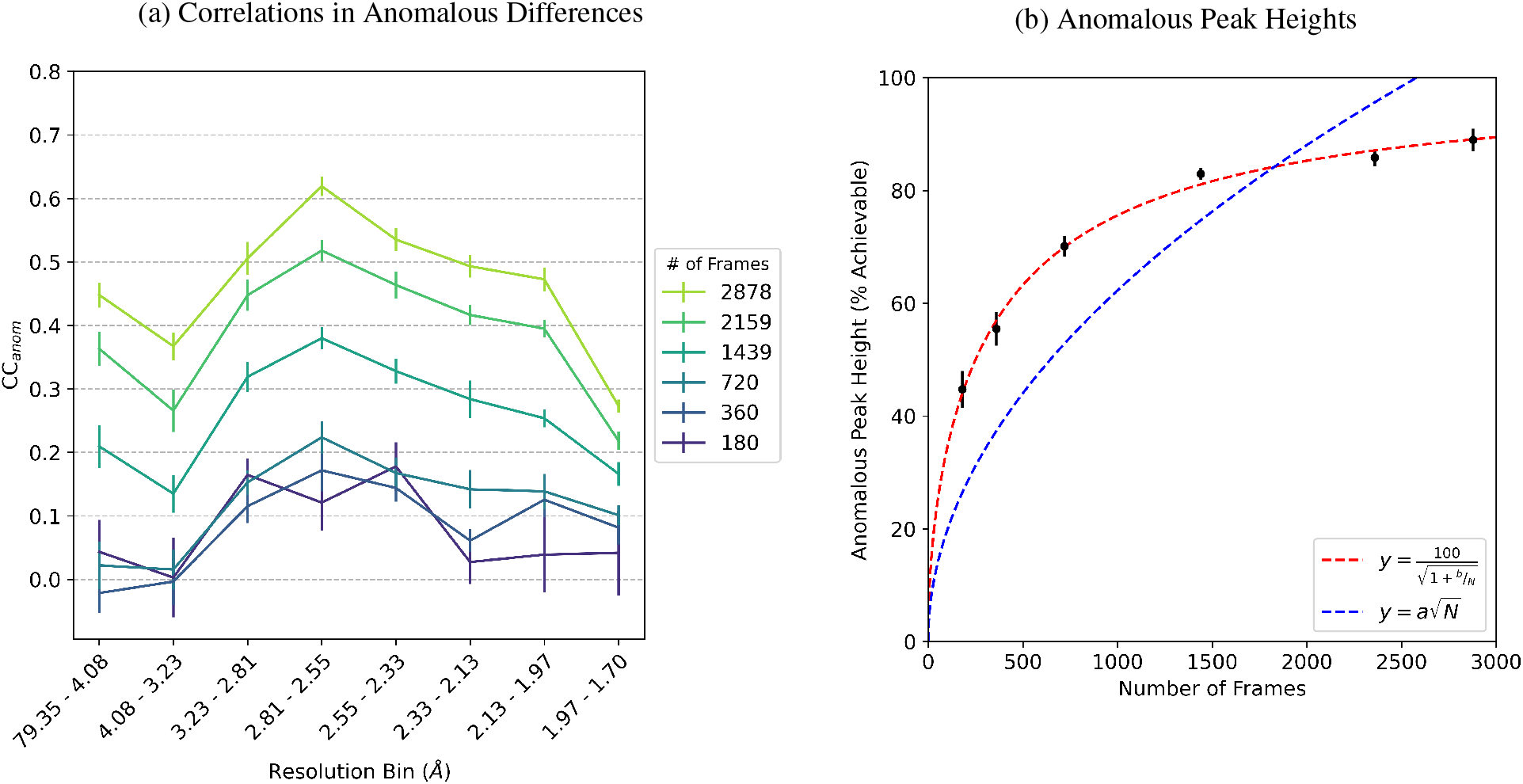
Redundancy improves the anomalous signal for a HEWL sulfur SAD dataset. (a) Spearman correlation coefficient for anomalous differences, *CC*_*anom*_, from repeated two-fold cross-validation for datasets composed of increasing numbers of frames. Error bars denote the mean ± standard deviation for 10 random partitions of the frames. The mean anomalous peak heights for the 10 sulfur atoms in the asymmetric unit cell after refinement using merged intensities for datasets with increasing numbers of frames (black; mean ± standard deviation). The peak heights are on an absolute scale relative to the maximal value that can be achieved. Non-linear least squares fits to the data are shown for 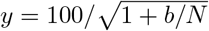 (red dashed line; *b* = 751 ± 75.8; 95% confidence interval) and 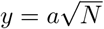 (blue dashed line; *a* = 1.97 ± 0.37; 95% confidence interval).

Since anomalous peak height relates directly to the correlation in measured anomalous differences across half-datasets [45], our extension of the theory in ref. [45] should also apply to predictions of the likelihood of successful phasing with the SAD method [46]. Consistent with this, we were only able to solve the HEWL structure by the SAD method with a dataset composed of at least 720 frames. Furthermore, the figures of merit of the solutions improved monotonically with increasing numbers of merged frames, and therefore with increasing multiplicity of observations (Table S1).

## 4 Discussion

Native SAD phasing relies on the accurate measurement of the small differences in intensity that arise due to anomalous scattering. As such, native SAD phasing can provide a rigorous test of the accuracy of new data collection approaches. Here, we presented a collection strategy at room temperature that takes advantage of the capabilities of modern macromolecular beamlines to collect high-quality, high-redundancy X-ray diffraction datasets. In particular, through the use of high attenuation and helical data collection, we were able to minimize the X-ray dose while taking advantage of the photon-counting, low-noise readout of the pixel array detectors that are in widespread use at macromolecular beamlines at synchrotrons.

To demonstrate this strategy, we used native anomalous signal to solve four structures of three model enzymes. The resulting anomalous difference maps clearly identify the sites of each anomalous scattering atom (Fig. 1). It was possible to observe anomalous scattering from manganese, phosphorous, and sulfur, and to resolve partial-occupancy sites for manganese in the DHFR structure that were not evident in the 2*F*_*o*_ − *F*_*c*_ maps (Fig. 2b). Furthermore, these room temperature structures contain significant conformational heterogeneity, as illustrated by the superposition of loop states observed in PTP1B (Fig. 3). These features emphasize the accuracy and sensitivity of the data collection approach presented here.

Although this work demonstrated native SAD phasing at 295 K using model enzymes with previously existing structures, this collection strategy could be used to solve new structures through experimental phasing or to collect high-quality datasets from single crystals near physiological temperatures. Although the crystals used in this project were fairly large (0.2-0.5 mm), the successful phasing of apo PTP1B required the merging of datasets from two different crystals. This illustrates that it is possible to use multi-crystal approaches, which may be an appealing way to achieve high-redundancy measurements due to the high isomorphism of protein crystals at room-temperature [13]. This could be an important application because multi-crystal approaches have been successful for native SAD phasing from crystals that diffract to more moderate resolutions (2.3-2.8 Å) [6].

Moreover, we extended the analysis of anomalous signal to explicitly account for the redundancy of observations. We found that higher redundancy of observed reflections improved the precision of anomalous differences (Fig. 4a), as well as the real-space anomalous signal (Fig. 4b), and that our model provides an excellent fit to the anomalous signal in all four datasets presented in this work (Fig. S2). For a given crystal with a particular resolution of diffraction, the observed anomalous signal improves with redundancy; however, this model demonstrates that real-space anomalous signal saturates at a map level that is dictated by the parameters of the diffraction experiment.

By demonstrating native SAD phasing at ambient temperature from single-crystal measurements, this work confirms that X-ray diffraction experiments at room temperature can produce highly accurate results that are not hindered by radiation damage. Since the high-redundancy strategy presented here can be implemented at many macromolecular beamlines, this work will support new investigations of the conformational ensembles of macromolecules near physiological temperatures.

## 5 Data Availability

All diffraction images used in this work have been deposited to SBGrid, with separate depositions for each structure [47, 48, 49, 50]. These depositions contain the complete processing details for DIALS, including the pixel mask that was used to exclude the beam stop shadow on the detector. The integrated intensities from DIALS and the scaled merged/unmerged data from AIMLESS have been uploaded to Zenodo for each image set [51, 52, 53, 54]. The refined structures presented here have been deposited with the Protein Data Bank [55] at the following PDB IDs: 7L84, 7LVC, 7MM1, and 7RIN.

## 6 Acknowledgements

We thank the staff at the Northeastern Collaborative Access Team (NE-CAT), beamline 24-ID-C of the Advanced Photon Source, for supporting our room-temperature crystallography experiments, with special thanks to Igor Kourinov for his assistance during data collection. We also thank Rachelle Gaudet (Harvard University) for helpful feedback on this manuscript.

## 8 Appendix

### 8.1 Impact of redundancy on anomalous signal

Terwilliger *et al* derived an expression relating the expected anomalous difference map peak height for a SAD experiment, 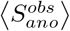, to the useful anomalous correlation of the dataset, *CC*_*ano*_ [45]:

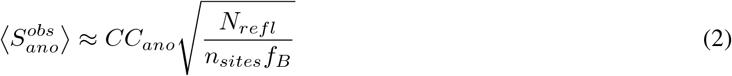

where *N*_*refl*_ is the number of observed reflections in the reciprocal asymmetric unit cell, *n*_*sites*_ is the number of sites in the anomalous substructure, and *f*_*B*_ is an average B-factor-modified anomalous atomic scattering factor. We build on their work to include the dependence of expected anomalous peak heights on the average redundancy of observations, *m*. To do that, we determine how *CC*_*ano*_ depends on *m*. Terwilliger *et al* also derived an expression for *CC*_*ano*_ in terms of a normalized variance, *E*^2^[45]:

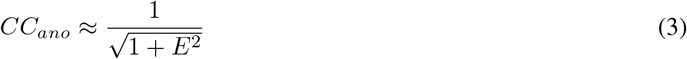

*E*^2^ is in turn defined as the ratio of the sum of the variance of the anomalous contribution that is not part of the anomalous substructure, 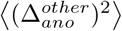, and the errors in measurement, *ε*^2^, to the variance in the useful anomalous signal, 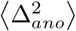 [45]:

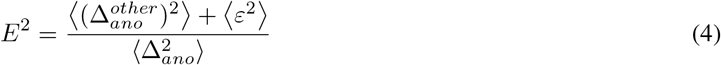

The measurement errors can be split up into fixed or systematic components, 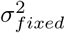, which do not improve with dataset size (e.g. due to phase errors and limitations of data processing), and random components, 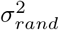, which do improve with multiplicity, *m*. If we assume that the random errors are uncorrelated, we can modify equation 4 as follows:

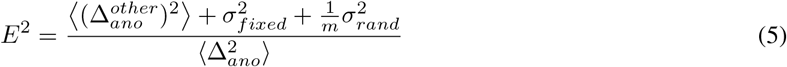

By substituting this expression into the equation for *CC*_*ano*_ from Terwilliger *et al*, we can modify equation 3 to become:

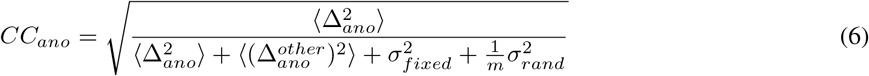

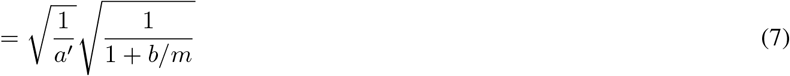

with 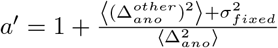 and 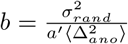.

Combining this expression for *CC*_*ano*_ with equation 2, we can relate the expected anomalous signal to the average redundancy of the observed reflections:

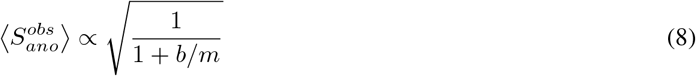

A practical implication of equation 8 is that anomalous peak height saturates as a function of multiplicity. However, for a given crystal that diffracts to a particular resolution increasing the multiplicity of the observations can decrease the random errors in the data, improving the anomalous signal.

## 9 Supplementary Figures

**Figure S1:**
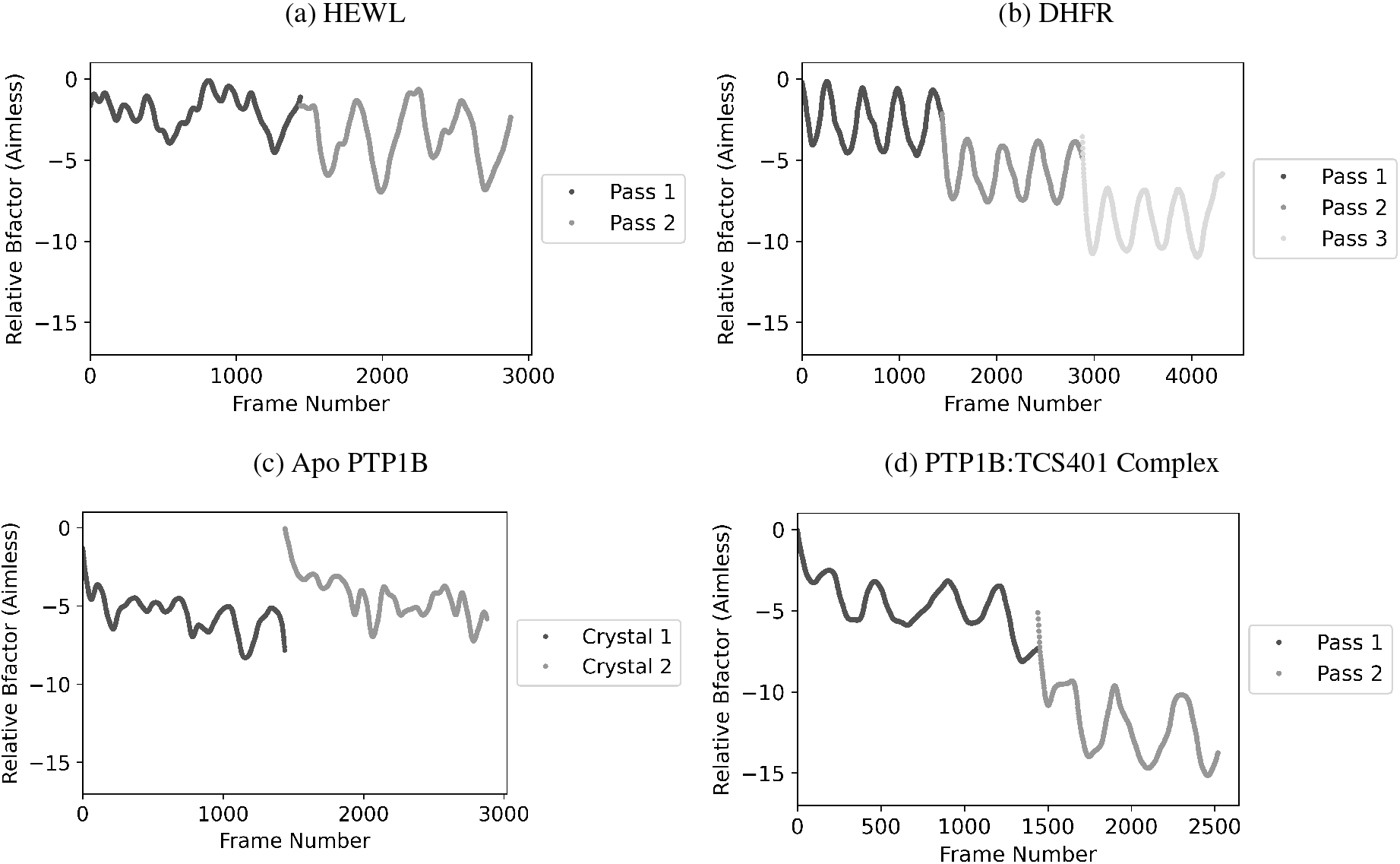
Relative image B-factors emphasize limited radiation damage. Image B-factors fit during scaling in *AIMLESS* are an indicator of radiation damage. Values significantly below -10 are often considered a sign of damage [25]. The image B-factors are presented for the datasets collected on each crystal for (a) HEWL, (b) DHFR, (c) apo PTP1B, and (d) PTP1B:TCS401 complex. Although the image B-factors for the final passes on DHFR and PTP1B:TCS401 reach values lower than -10, excluding such frames from the analysis was not found to improve the SAD phasing results (not shown).

**Figure S2:**
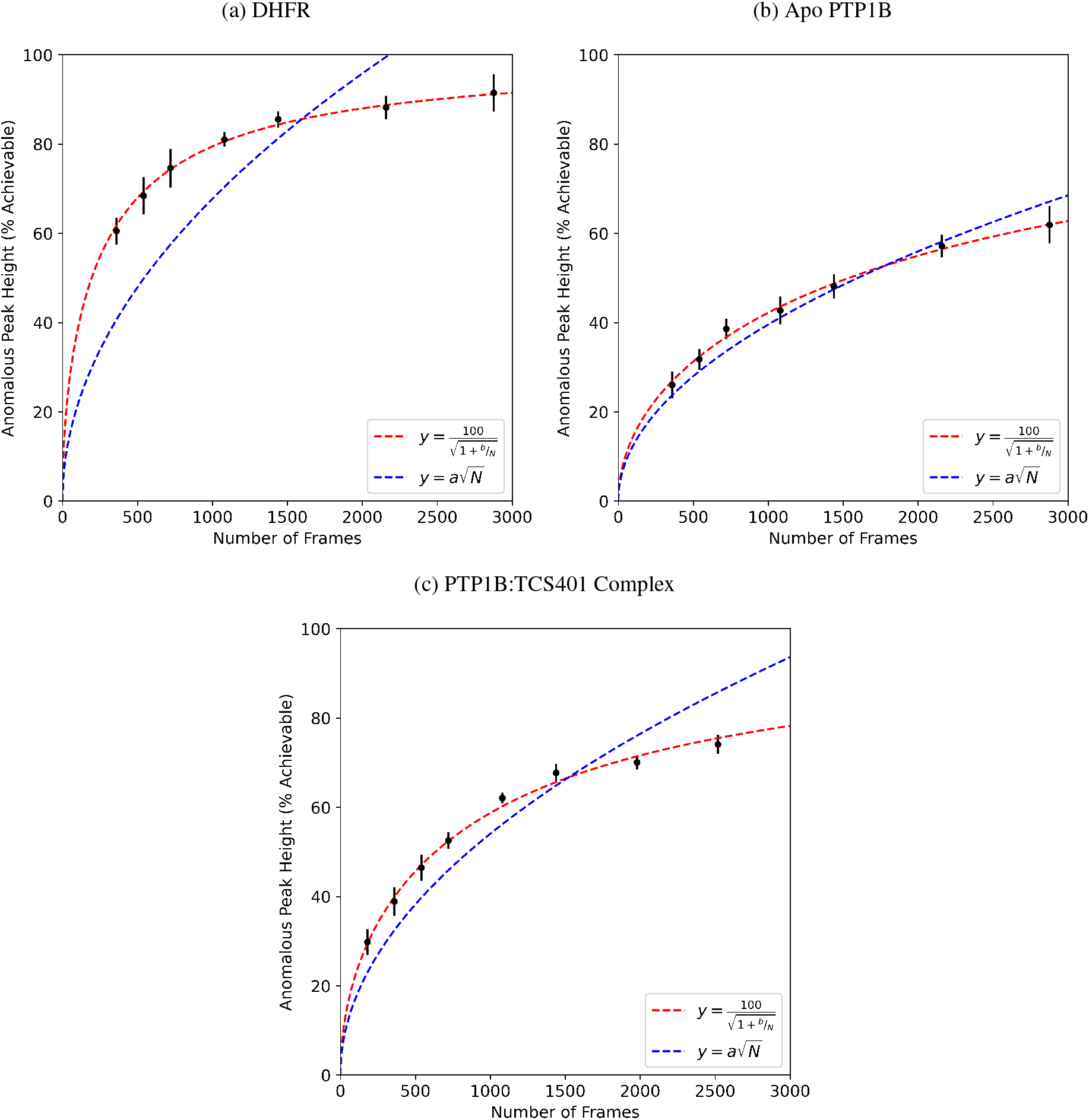
Redundancy improves anomalous signal for native SAD datasets. Anomalous peak heights corresponding to the anomalous sites in the asymmetric unit cell after refinement using merged intensities for datasets with different numbers of frames are shown for (a) DHFR, (b) Apo PTP1B, and (c) PTP1B:TCS401 complex (black data points; mean ± standard deviation). The peak heights are on an absolute scale relative to the maximal value that can be achieved. Non-linear least squares fits to the data are shown for 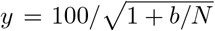 (red dashed line) and 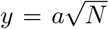 (blue dashed line). This analysis only includes the anomalous sites that could be identified in all anomalous difference maps for the subsets of the data. The values fit for *b* are (a) 583 ± 48.1, (b) 4620 ± 1140, and (c) 1900 ± 250 (95% confidence interval). The values fit for *a* are (a) 2.14 ± 0.36, (b) 1.25 ± 0.07, and (c) 1.71 ± 0.16 (95% confidence interval).

**Table S1:** Figure of Merit from Native SAD Phasing of HEWL

| **Number of Frames** | **Figure of Merit** |
| --- | --- |
| 180 | 0.23 |
| 360 | 0.24 |
| 720 | 0.40 |
| 1439 | 0.44 |
| 2159 | 0.46 |
| 2878 | 0.47 |

